# Coupling Step-Wise Motility and Traction Force Patterns in chemotaxing cells

**DOI:** 10.1101/2025.10.28.685178

**Authors:** Sebastián Echeverría-Alar, Wouter-Jan Rappel

## Abstract

Chemotaxing Dictyostelium discoideum cells migrate in a step-wise fashion characterized by periodic protrusion, contraction, and rear retraction cycles accompanied by distinct traction force patterns. Traction force microscopy reveals two stationary force spots that exchange identity as the cell advances and generate a convergent stress pattern with both axial and lateral components. To investigate the physical origin of these traction patterns, we developed a continuum, phase-field model that couples cytosolic flow, active stresses, and substrate friction within a cell with a deformable morpholgy. The model incorporates protrusive forces at the front, contractile stresses at the rear and sides, and spatially localized adhesive regions that undergo cyclic activation. While this baseline model reproduces persistent motion, it fails to capture the experimentally observed traction force patterns and cell morphology. Guided by new experiments visualizing myosin dynamics, we extended the model to include a localized contractile myosin patch positioned between the two adhesive regions. This modification yields cell shapes, speeds, and convergent traction patterns consistent with experimental measurements. The results demonstrate that a centrally positioned myosin patch is sufficient to generate the step-wise migration cycle and the characteristic convergent traction pattern of Dictyostelium cells, providing a mechanistic link between intracellular contractility, cytosolic flow, and force transmission during amoeboid motility.

## I. INTRODUCTION

Amoeboid motility is central to processes ranging from embryonic morphogenesis to immune surveillance [1]. One of the most-used model systems to investigate this motility is the soil amoeba *Dictyostelium discoideum* (Dicty for short) [2, 3]. Dicty cells move fast compared to other cell types (roughly one cell diameter ( ≈ 10 *µ*m) per minute), do not require temperature regulation, and can be easily genetically modified to create mutants and fluorescent reporters [4]. Furthermore, once deprived of food, these cells chemotax up gradients of the small molecule cAMP, offering experimentalists ways to steer and direct their motion [5–7].

Experiments have shown that chemotaxing Dicty cells undergo quasiperiodic motility cycles consisting of protrusion, contraction, rear retraction, and relaxation, accompanied by oscillations in cell length [8, 9]. During this motion, cells exhibit a characteristic elongation along the axis of migration, tapering off in the back. These experiments also quantified cell–substrate forces using traction force microscopy, which computes traction stresses by measuring cell-induced displacements of embedded beads in compliant gels [10], revealing oscillations in strain energy and substrate stresses. A striking feature of these data is a step-wise mode of movement in which inward-directed traction adhesions remain nearly stationary in the lab frame as the cell advances and “steps” from old to newly formed anterior sites. In addition to these axial (anterior–posterior) forces, experiments also showed the importance of lateral squeezing. Together, axial and lateral tensions produce characteristic kymographs that reveal traction force spots at the front and rear during persistent migration, while the cell maintains an elongated shape.

Despite these detailed measurements, the physical origin of the observed traction patterns remains incompletely understood. Existing interpretations emphasize actin polymerization at the front, myosin-II–driven contractility at the rear, and adhesion turnover, but they do not yet explain how cell movement gives rise to the observed traction force patterns [11]. One possibility is that intracellular flows couple to the substrate to generate the specific spatiotemporal force signatures during the stepping cycle. These flows are thought to be especially important in the case of Dicty since, in contrast to most mammalian cells, the adhesion between cells and the substrate are not dependent on focal adhesion complexes or integrins. Instead, the adhesive forces between Dicty cells and the substrate are believed to be non-specific, resulting in the ability of Dicty cells to move on a wide variety of surfaces [12–14].

Here, we investigated possible mechanisms that can explain the observed traction force patterns, using continuum, physics-based models. In these models, we represented the cell with a phase-field description of a moving, deformable boundary. This description is particularly well-suited to model cells with deformable morphologies as it does not require explicit tracking of the interface [15–19]. Cell deformations can be coupled to a viscous cytoplasm such that active stresses and pressure gradients drive cytosolic flow [20]. The resulting shear of the cytosolic flow against the substrate yields a distributed frictional traction whose axial and lateral components are determined by the evolving cell shape and internal hydrodynamics. In our model, a chemotactic gradient provided a driving force such that the cell moves in a direct fashion. Furthermore, we included constant lateral forces and also introduced spatially localized adhesive regions that exchange identity during each migration cycle (i.e., the anterior region becomes posterior).

We found that this model was not able to reproduce both the experimental traction force patterns and the elongated cell shape. Motivated by this discrepancy, we carried out experiments in which we visualized myosin, the motor protein responsible for cell contractions, in chemotaxing cells. These experiments revealed the existence of a myosin patch between the two traction force spots, stationary in the laboratory frame of reference. When we implemented this myosin patch into the model, we found cytosolic flow patterns, cell speeds, and average cell shapes that were consistent with experiments. Thus, our study offers a mechanistic explanation for the stepping mode of amoeboid migration, which relies on an overlooked role of myosin.

This paper is organized as follows: in Section II we recapitulate the previous experimental results [9], and in Section III we describe and apply our phase-field model and attempt to reproduce the experimental results. In section 4, we describe new experimental results, include a novel myosin distribution, while in section 5 we incorporate this distribution and show that the extended model can produce results that are consistent with the experiments. Finally, we conclude with a discussion.

## II. EXPERIMENTAL RECAP

Using traction force microscopy on compliant polyacrylamide gels, Bastounis *et al*. quantified the spatiotemporal pattern of forces generated by chemotaxing Dicty cells over many migration cycles (schematically shown in Fig. 1. [9]. Cells moved with a speed of approximately 10 *µ*m/min and kymographs aligned to the cell’s anterior–posterior axis revealed a robust, step-like motion: two spatially localized axial traction regions persist stationary in the lab frame as the cell advances. During each cycle, a new anterior traction region forms ahead of the centroid, the previous anterior region transitions to the rear, and the former posterior region weakens, producing a front–back “exchange of identity.” These force cycles are accompanied by a distinct average morphological shape, consisting of a broader front and a narrower back (Fig. 1). These experiments also revealed a characteristic convergent traction force pattern, consisting on strong axial forces, directed toward the center of the cell, together with weaker lateral forces that “squeeze” the cell and are also directed towards the cell’s center.

**FIG. 1.**
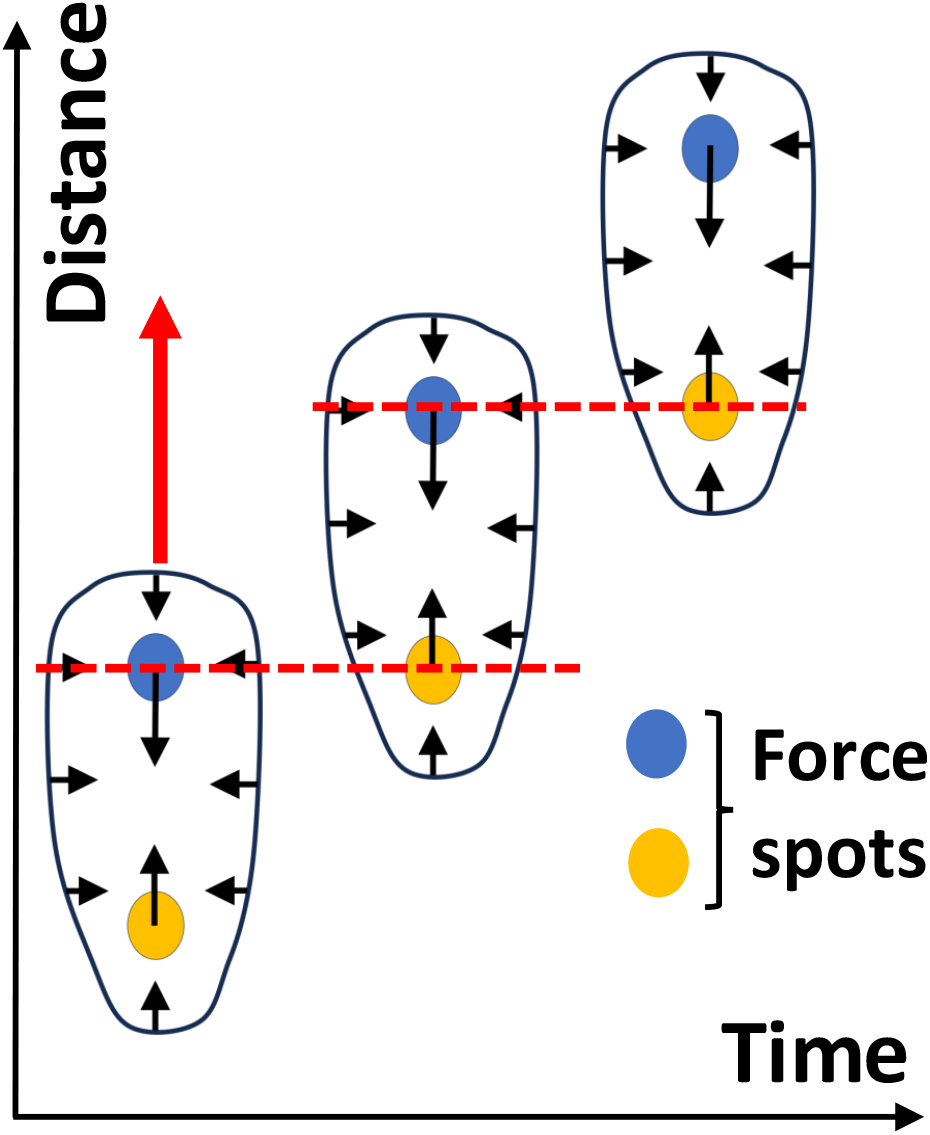
Schematic representation of a chemotaxing Dicty cells, shown at three different time points and moving in a step-wise fashion as indicated by the red arrow. Traction forces are concentrated at the two force spots (blue and yellow disks) and the time-averaged shape is broader at the front than at the back. The force spots remain stationary in the laboratory frame and the anterior spot becomes the posterior one as the cell migrates forward (schematically indicated by the red dashed line). The resulting experimentally observed traction force pattern is indicated by the black arrows and shows a distinct convergent pattern: both axial and lateral forces are directed towards the center of the cell.

## III. INITIAL MODEL

We first determined whether a model that incorporates adhesive spots together with lateral adhesion, area conservation, cytosolic flow, and protrusive and contractile forces at the front, back and sides, respectively, can reproduce the experimental results. We used the phase field approach, in which a scalar field *φ*(**r**, *t*) provides a smooth and continuous representation of the cell. Specifically, *φ* = 1 inside the cell and *φ* = 0 outside the cell, with the two states connected through an interface of width *ϵ*. This parameter was fixed to *ϵ* = 2 *µ*m (roughly 10Δ*x*) to ensure smoothness and numerical stability of the phase field dynamics. The spatiotemporal evolution of the cell membrane was given by

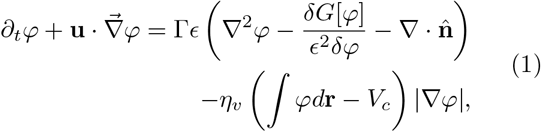

where the cell membrane was advected by the cytoskeletal flow with velocity **u**. The first term on the righthand side of Eq. (1) was added for stability purposes [20, 21], imposing a relaxational dynamics controlled by the constant Γ. Here, *G*(*φ*) = 18*φ*^2^(1 − *φ*)^2^ is a double well potential with minima at *φ* = 0 and *φ* = 1, and 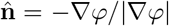 is the unit normal vector. The last term, proportional to *η*_*v*_, penalizes deviations from the prescribed cell size *V*_*c*_.

Following previous studies, we modeled the cytoskeleton of the cell as an active, viscous and compressible fluid [20, 22, 23]. In this framework, the cytoskeletal flow can be described with a Stokes equation representing a momentum balance in the inertialess limit:

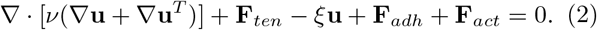

The first term in this equation corresponds to viscous stresses, parameterized by the viscosity coefficient of the cytosol *ν* [24–26], while the second term represents the membrane tension. In equilibrium, this tension is associated with a Ginzburg-Landau type of energy ℱ [*φ*, ∇*φ*] = *γ ∫* (*ϵ* | ∇ *φ*| ^2^*/*2 + *G*(*φ*)*/ϵ*)*d***r**, such that the force can be expressed as 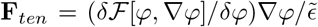, where 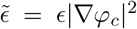 and *γ* denotes the surface tension [16]. The third term models cell-substrate friction, using a simple linear drag law with coefficient *ξ* [24].

The fourth term in Eq. (2) accounts for adhesive forces and comprises two components: (i) adhesive complexes associated with the stepping motion along the direction of migration 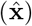, **F**_*ss*_, and (ii) lateral forces **F**_*lat*_, which stabilize directed motion, **F**_*adh*_ = **F**_*ss*_ + **F**_*lat*_. To model the adhesive complexes, we define a series of *N* equidistant circular regions along the horizontal axis of the computational box 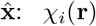 (Fig. 2), each with a time-dependent strength *η*_*i*_(*t*) (*i* = 1,.., *N* ). The resulting traction force reads

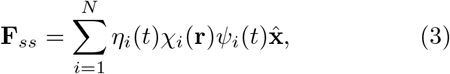

where *ψ*(*t*) indicates whether the adhesive complex *i* is active (*ψ* = 1) or inactive (*ψ* = 0). Activation of a front complex occurs when the cell fully covers the adhesive region. Inspired by experimental observations, we imposed a simple switching rule dynamics for the stepping dynamics. The initial configuration was a cell with two active adhesive complexes: a front (*i* = *f* (*t*)) and a back (*i* = *b*(*t*)), ∑*ψ*_*i*_(*t*) = 2. As the cell migrated to the right and eventually a new adhesive complex became active, ∑ *ψ*_*i*_(*t*) = 3, the front and back were updated as

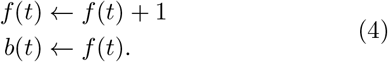

**FIG. 2.**
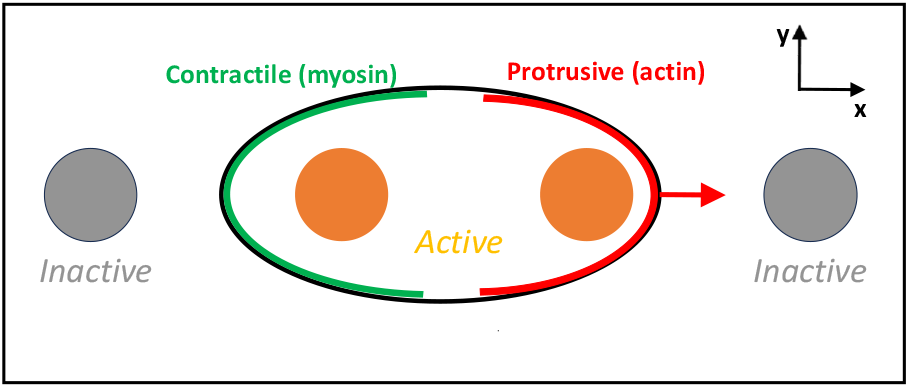
Model set-up. A computational domain contains four equally-spaced adhesive spots. Only the spots that are within the computational cell are active (orange) while the ones outside are inactive (gray). Protrusive forces (red) are localized at the front of the cell while contractile forces (green) are present at the back of the cell. Cell is migrating in the **x** direction indicated by the arrow.

The previous back remained active until the cell no longer covered it [9], returning to ∑ *ψ*_*i*_(*t*) = 2. This stepping cycle repeated periodically and adhesive strengths were given by

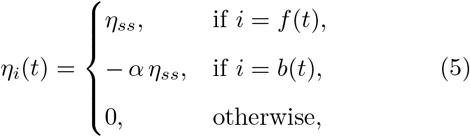

with *α <* 1 and *η*_*ss*_ constants. The lateral traction forces were localized at the cell interface by making it proportional to the derivative of the phase field:

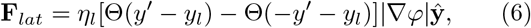

where *y*^*′*^ = *y* − *y*_*c*_, with *y*_*c*_ the *y*-coordinate of the cell’s center of mass, and *y*_*l*_ controls the spatial extent of the force. In this study, we have chosen *N* = 4, which ensures that there is always at least one inactive adhesive spot.

The active forces, **F**_*act*_, were dictated by active stresses driving cytoskeletal flow, which depend on the spatial distribution of F-actin and myosin. The F-actin distribution followed the classical observation in migratory cells, in which actin polymerization occurs at the cell front. The corresponding active stress, which promotes protrusion growth, was modeled as

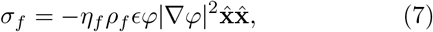

where *ρ*_*f*_ = Θ(*x* − *x*_*c*_) is the F-actin density, *x*_*c*_ is the *x*-coordinate of the cell’s center of mass, and *η*_*f*_ sets the strength of the stress. Unlike previous studies [20], the protrusion growth was biased along the direction of migration 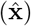, mimicking a cell sensing a chemoattractant gradient to the right.

The myosin-driven contractility in this model was divided into two contributions. The first contribution was inspired by the ellipsoidal shapes exhibited by fast-migrating cells [9, 27]. We hypothesized that a contractile stress acting along the direction orthogonal to migration (**ŷ**), possibly triggered by a strong chemoattractant gradient, favors the ellipsoidal morphology. This stress was modeled as

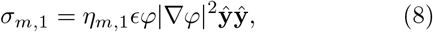

where *η*_*m*,1_ controls the stress magnitude. The second contribution was introduced to contract the cell back and generate effective forward motion. We assumed that a localized myosin distribution *ρ*_*m*,2_ along the cell boundary produces both inward and tangential contractile stresses

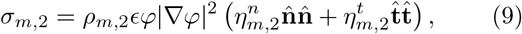

where *ρ*_*m*,2_ = Θ(*x*_*m*2_ − *x*^*′*^), with *x*_*m*2_ a spatial coordinate defining the region of myosin localization. The parameters 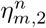 and 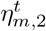 set the strengths of the normal and tangential stresses, respectively, and 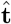 is the unit tangential vector (a 90^*°*^ counterclockwise rotation of 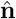 ). Finally, the active force was obtained as

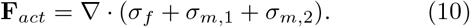

The model was simulated on a *m* × *m* square (here taken to be *m* = 168) computational domain with periodic boundary conditions along the horizontal direction and with a grid spacing of Δ *x* = 0.19 *µ*m. As initial condition, we chose an elliptical cell (Fig. 2). The equations were solved as detailed earlier [23]: the phase field equation was time-stepped using a forward Euler scheme (Δ *t* = 0.002 s) while the Stokes equation was solved with a semi-implicit Fourier spectral scheme. The latter was iterated until the difference between the velocity field at the *n*^*th*^ + 1 and *n*^*th*^ timestep became smaller than a prescribed value (max |*u*^(*n*+1)^ − *u*^(*n*)^ | *<* 0.01 max *u*^(*n*)^ | ) or when 50 iterations were completed.

Solving the model equations resulted in a cell moving to the right, accompanied by a cytosolic flow pattern that generated traction forces. To determine if the presence of the adhesive spots, together with the incorporated active forces, were able to produce traction force patterns, morphologies, and cell speeds consistent with the experiments, we systematically varied force and adhesive parameters. Specifically, we varied parameters { *η*_*f*_, *η*_*m*,1_, *η*_*m*,2_, *η*_*l*_, *α* }, and measured the average speed *V* along with the shape ratios between the cell length in the direction of motion and the widths of the front (*R*_*f*_ ) and back (*R*_*b*_) (Fig. 3). The average speed was obtained from cell migration simulations over a time window of 500 s. *R*_*f*_ was measured 5*ϵ* to the left of the cell’s front, while *R*_*b*_ was measured 5*ϵ* to the right of the cell’s back. Changing of parameters revealed that the model was not able to replicate the experimental results: *V* within the range 10 − 15 *µ*m/min, *R*_*b*_ ≈ 3 and *R*_*f*_ ≈ 4 [9]. In all parameter combinations, we did not observe speeds exceeding 10 *µ*m/min, and in most cases we found aberrant morphologies in which the front was narrower than the back. Furthermore, even though the traction force patterns were always convergent, the lateral (*y* − ) component was consistently too small.

**FIG. 3.**
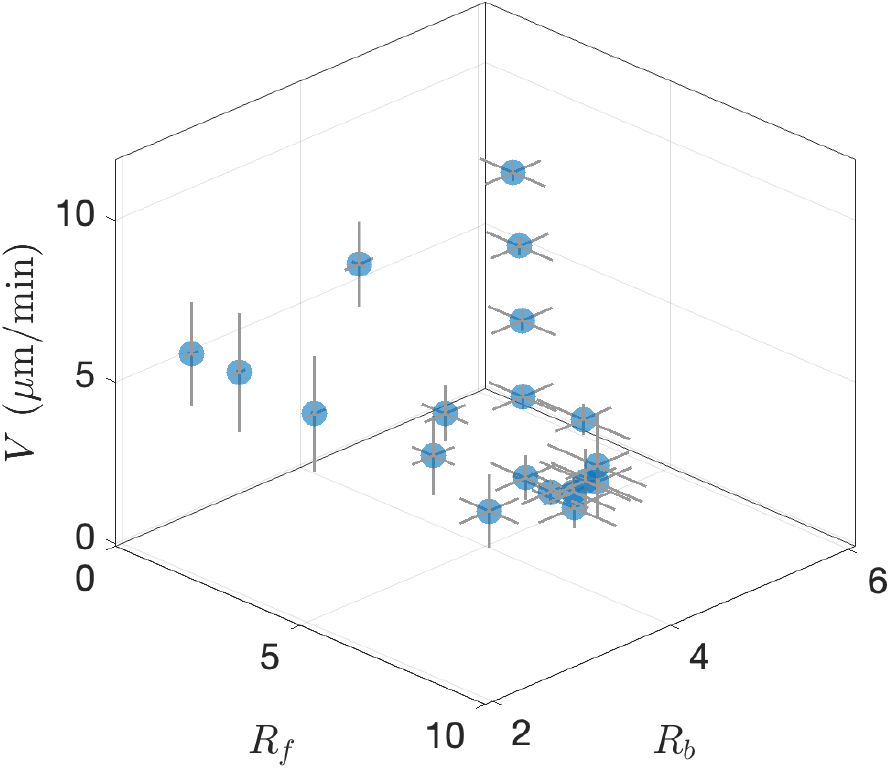
Speed and morphological measurements for different combination of the parameters { *η*_*f*_, *η*_*m*,1_, *η*_*m*,2_, *η*_*l*_, *α* }. The bars indicate standard deviations, obtained by following the cell for a prolonged period of time.

## IV. MODEL EXTENSION

Motivated by our simulation results and hypothesizing that molecular distributions can play an important role in generating traction forces, we carried out traction force experiments using Dicty cells in which myosin is fluorescently tagged. In these experiments, traction force microscopy and imaging were carried out as previously detailed and used wild-type AX2 cells, transformed with the plasmid expressing pBig-myo, expressing GFP-myoII [23]. Cells were plated on a deformable substrate that contained small fluorescent tracer particles. Fluorescent images (561nm excitation, for the red fluorescent beads, and 488nm for the GFP probe) were captured every 15 s with a 63X oil objective on a spinning-disk confocal Zeiss Axio Observer inverted microscope equipped with a Roper Quantum 512SC camera. Autofocus was set on the fluorescent beads at the surface of the gel so that all images were recorded in the basal plane. The spatial map of displacements of tracer particles (relative to their positions with no cells on the substrate) was measured and converted, using an open source MATLAB algorithm (2018a; The Mathworks) [28], to a spatial map of cell traction forces. Recordings started 15 min after plating for up to 3 h. For ease of visualization, stress maps and fluorescent images were rotated as described before [23].

As in the original study, these chemotaxing cells were found to be moving in a step-wise fashion with distinct anterior/posterior force spots that remained stationary and are cyclically exchanged. An example of a kymograph is shown in Fig. 4A, where the traction forces are visualized in blue and where the red dashed lines highlight the front/back exchange of the traction force spots. However, these experiments also revealed a distinct myosin patch, located in between the two stationary spots (green signal in Fig. 4A).

**FIG. 4.**
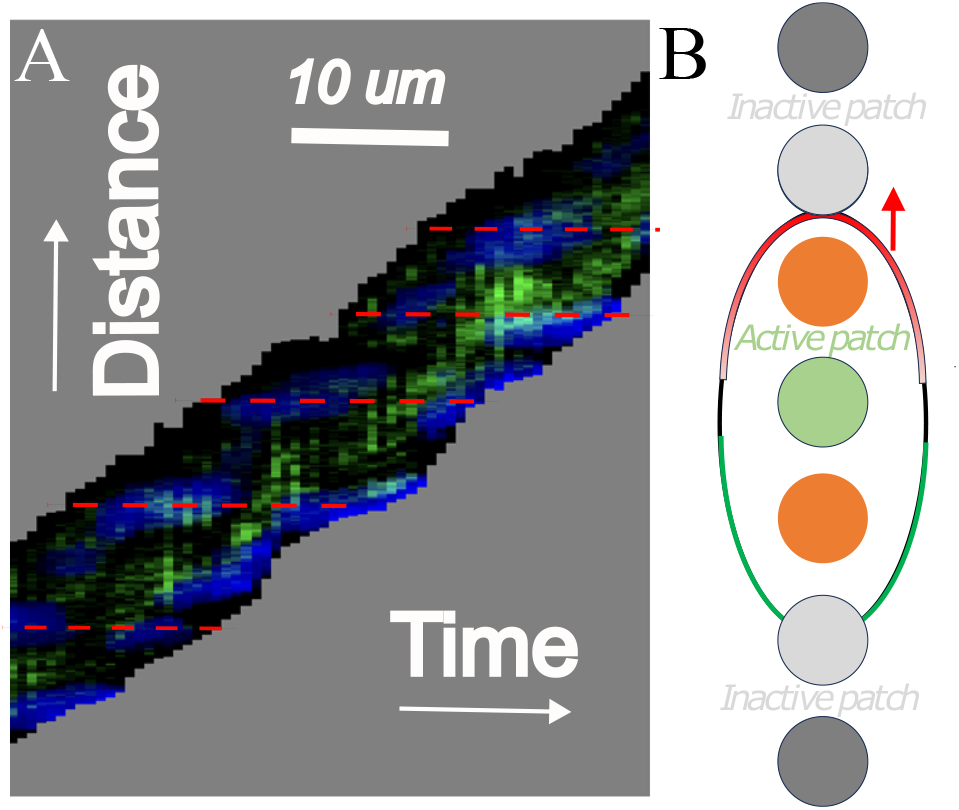
(A) kymograph of a chemotaxing Dicty cell with traction stress visualized in blue and myosin in green. The adhesive spots remain stationary in the laboratory frame, as indicated by the red dashed line, while myosin is mainly present along the length axis and in between the adhesive spots. (B) Schematic drawing of the new model set-up with an active contractile myosin patch (green) located between two active adhesive complexes (orange). Inactive myosin (light gray) and inactive adhesive patches (dark gray) are also shown. Red arrow indicated direction of motion.

Next, we incorporated these myosin patches into our model and assumed that they connect adjacent adhesive complexes involved in the stepping motion (Fig. 4B). We defined four myosin patches centered between adjacent adhesive complexes. Each patch has a Gaussian-distributed myosin profile to avoid singularities when computing forces. Thus, the total myosin density of this new contribution is

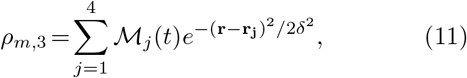

where **r**_**j**_ is the center of patch *j* and *δ* is the Gaussian width. The variable ℳ_*j*_ is equal to 1 if and only if the adjacent adhesive complexes are active (*ψ* = 1). The corresponding isotropic contractile stress is

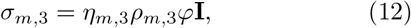

parameterized by *η*_*m*,3_. As a result of this contribution, the active force now reads

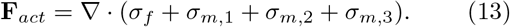

We incorporated this experimentally motivated contractile spot into the model and systematically explored parameter space, which now included the strength of this spot (*η*_*m*,3_) and its size *δ*. The extended model recapitulated key features of chemotaxing cells, including speed, shape, and convergent traction patterns. In particular, we found that achieving a front broader than the rear and speeds consistent with experiments required both a minimum contractile strength and patch size for the myosin domains. This is shown in the phase diagram spanning the strength-size space where we plot the cell speed using a color map (Fig. 5A). Along the diagonal of this diagram, indicated by the star symbols, the speeds were consistent with the experiments (*V* ∼10µm/min). The cell morphology for these parameter combinations was also consistent with the experimental shape, showing a cell with a broader front and a narrower back. Furthermore, the traction force pattern was convergent and showed large axial forces directed toward the cell centroid, together with smaller lateral forces (upper panel in Fig. 5).

**FIG. 5.**
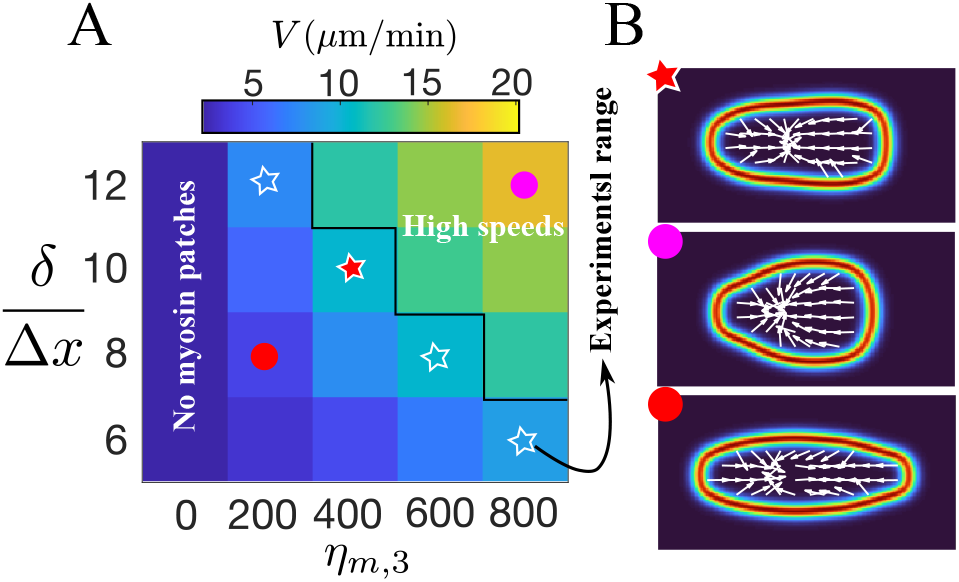
Inclusion of a contractile myosin patch in cell simulations. (A) Speed of the cell’s centroid, *V*, in the parameter space spanned by the contractile strength of the patch, *η*_*m*,3_, and its size,*δ*, normalized by the spatial discretization. Speeds within the experimental range are indicated with the star symbols. (B) Three cell shapes with their respective traction patterns corresponding to different migrating speeds: within the experimental range (top panel), above the experimental range (middle panel), and below the experimental range (bottom panel).

Our simulations showed that cells with parameter value combinations above this diagonal migrated faster, with speeds in the range of 15 − 21 *µ*m/min. These cells, however, displayed a markedly different morphology, with a much wider front and a less elongated shape (middle panel in Fig. 5). By contrast, for parameter combinations below the diagonal, cells were significantly slower (*v <* 10*µ*m/min) and exhibited a larger *R*_*b*_ than *R*_*f*_ (bottom panel in Fig. 5). Thus, parameter combinations away from the diagonal produced results that were inconsistent with experiments, suggesting that Dicty cells migrate with patch size and strength combinations that are correlated.

## V. DISCUSSION

Our modeling results highlight the importance of molecular distributions in creating physiologically relevant mixtures of cell speed, cell shape and traction force patterns. A model in which contractile forces are only concentrated at the back and side of a migrating cell was not able to reproduce experimental results. Only when we added an additional contractile force, arising from a myosin patch that was centered between the two adhesive spots, were we able to create results that are consistent with experiments. It is worth pointing out that our study has several limitations. First, our model does not include pseudopod formation, and can thus only be compared to average shapes in the experiment. Second, we do not include a mechanism for the creation of the alternating adhesive spots. Further work is needed to determine the molecular nature of these spots and how they may be formed. Nevertheless, our model points towards the importance of molecular distributions within the cell. It would be interesting to see if other cell types that also show the step by step motion (for example, HL60 cells) show a similar myosin patch. We are currently exploring this possibility.

## ACKNOWLEDGMENTS

This work was supported by NSF MCB 2426002 and NSF PHY 2310496.

